# Provision of essential resources as a persistence strategy in food webs

**DOI:** 10.1101/2023.01.27.525839

**Authors:** Michael Raatz

## Abstract

Pairwise interactions in food webs, including those between predator and prey are often modulated by a third species. Such higher-order interactions are important structural components of natural food webs that can increase the stability of communities against perturbations and ensure continued ecosystem functioning. Particularly the flux of rare organic and inorganic compounds that are essential to species in the community can create higher-order interactions. Even though many such compounds exist, their effect on structuring communities is little understood. In this study, I perform invasion analyses on a general food web model that depicts apparent and exploitative competition. Introducing the provision of essential resources by a prey species to either its competitor or its predator as a higher-order interaction, I find that this mechanism can ensure the focal prey’s persistence. Larger dietary essentiality, i.e. a stronger dependence of the predator or the competitor on the essential resource can increase the invasion growth rate of the focal prey to positive values, thus promoting its persistence when it would go extinct for low essentiality. This research shows that essential resources and the higher-order interactions created by them should be considered in community ecology.

## Introduction

Growth, reproduction and survival of organisms can be limited by organic and inorganic compounds, which often are not present in the organism’s diet in favourable concentrations or ratios. Particularly consumers at the plant-herbivore interface are often affected by dietary mismatches (Elser et al., 1996; Gaedke et al., 2002; Wacker & Martin-Creuzburg, 2012; Urabe et al., 2018). This motivated considering besides food quantity also the quality of food when investigating performance measures of aquatic consumers (Andersen et al., 2004; Anderson & Hessen, 2005; Wacker & Martin-Creuzburg, 2012; Guo et al., 2016; Schälicke et al., 2019; Koussoroplis et al., 2019), terrestrial herbivores (Douglas, 2015; Eberl et al., 2020) and pollinators (Filipiak et al., 2017). The scarcity of resources that are essential for growth and reproduction but cannot be easily acquired from the environment can constrain the flow of matter and energy between trophic levels. Therefore, dietary limitations induced by essential resources can have important effects on population and community dynamics (Muller et al., 2001; Gaedke et al., 2002; Schade et al., 2003; Stiefs et al., 2010; Iwabuchi & Urabe, 2012; Singer et al., 2012; Raatz et al., 2017; Burian et al., 2020).

Dietary dependencies also regularly exist within the same trophic level, where uptake of essential resources occurs from the environment, such as within the microbial loop when bacteria consume dissolved organic carbon from phytoplankton exudates (Azam et al., 1983; Pomeroy et al., 2007) or during the exchange of essential nutrients and metabolites between bacteria and microalgae (Soria-Dengg et al., 2001; Croft et al., 2005; Kazamia et al., 2012; Suleiman et al., 2016; D’Souza et al., 2018; Oña & Kost, 2022). Understanding the mechanisms and effects of such dependencies is crucial for biodiversity research given for example the importance of microalgae such as diatoms for aquatic ecosystems and global carbon dynamics (Croft et al., 2005; Koedooder et al., 2019), but also for medical fields like human microbiome research (Herren, 2020) and antibiotic resistance research (Adamowicz et al., 2018). Taken together, dietary mismatches and dependencies of organisms from the same or different trophic levels are crucial determinants for the structure of their communities. Mechanistically, community structure is determined by direct interactions within pairs of species or by indirect interactions across multiple species from the same or different trophic levels, e.g. through trophic cascades or apparent competition. Additionally to direct and indirect interactions, higher-order interactions, here defined as the density of a third species affecting the interaction of two other species (sensu Billick & Case, 1994), were found to potentially structure communities. The effects of higher-order interactions include stabilizing population dynamics (Grilli et al., 2017), increasing robustness against perturbation (Terry et al., 2019; Gibbs et al., 2023), determining fitness of competitors (Mayfield & Stouffer, 2017) and affecting biodiversity-ecosystem-functioning relationships (Miele et al., 2019). Examples for higher-order interactions include trait-mediated effects such as a predator affecting the foraging rate of its prey or the prey’s predation risk from other predators, and environment-mediated effects such as one species providing refuge to another species (Wootton, 2002; Miele et al., 2019).

In this paper, I will investigate another, so far unrecognized mechanism for creating higher-order interactions that arises from the provisioning of essential resources. In the presence of dietary mismatches one species, from here on referred to as the focal species, may provide resources that are essential to other community members. Such interactions are possible both towards members of the same trophic level, such as competitors, or towards members of different trophic levels, e.g. shared predators that prey on multiple species. For example, a higher-order interaction within the same trophic level is created when a competitor is co-limited by two resources but can only obtain one of those two resources from its environment and relies on another prey (the focal prey) to provide the other co-limiting resource. This provision may occur for example by leakage of common goods (Gore et al., 2009) or carbon exudation in otherwise carbon-limited environments (Bratbak & Thingstad, 1985; Raatz et al., 2018). A higher-order interaction between different trophic levels can arise when a predator obtains energy from multiple prey species but only the focal prey may provide additional, essential resources, e.g. vitamins or polyunsaturated fatty acids (Wacker & Martin-Creuzburg, 2012). Excess essential resources provided by the focal prey may then be used to efficiently convert other low-quality prey into predator biomass (Raatz et al., 2017).

In these two cases the provision of essential resources by the focal prey creates a higher-order interaction that manifests as an interaction modification (sensu Terry et al., 2019) that regulates the flow of matter to the competitor or predator compartment in these communities, respectively. Regulating such fluxes has the potential to affect the biomass distribution in the community, ultimately determining the persistence of individual species. If such higher-order interactions increase the persistence of the focal prey they pose as an example for a niche-improving form of niche construction and they may thus even be adaptive (Kylafis & Loreau, 2008, 2011; Laland et al., 2016). Consequently, in this paper, I will establish the provision of essential resources in a community as a mechanism driving higher-order interactions that may increase the persistence of the focal prey species and prevent its extinction either from predation or competition.

## Methods

Investigating persistence of a focal species typically employs invasion analysis, which determines the net growth rate of that species in the remaining resident community when it is rare (and assumed to be invading) (MacArthur & Levins, 1967; Chesson, 1994; Ellner et al., 2019). If the focal prey provides the essential resources to some components of the community, being rare equates to switching off the higher-order interaction. Invasion analysis is therefore the perfect tool for determining the effect of essential resources both on the resident community and the persistence of the focal prey. Accordingly, I will investigate the invasion growth rate of the focal prey species *X*_1_ in a community that contains an abiotic resource *R*, a competing species *X*_2_ and a shared predator *Y* (Eqn. 1, Fig. 1), to incorporate essential resource provision in food webs. I assume a chemostat-type model in which the abiotic resource *R* is provided at a constant rate *R*_0_ *δ* and all entities experience the same dilution, see Tab. 1 for parameter definitions and values. The two prey species *X*_1_ and *X*_2_ take up the abiotic resource *R* at some rate *r u*(*R*), where *u*(*R*) defines the functional form of prey resource uptake. Both prey species are consumed by the predator following a functional response *f* (*X*_1_*, X*_2_). I assume that the two prey species differ only in their attack probability (sensu Ehrlich & Gaedke, 2018) by a factor *p* and their maximum growth rate by a factor *ϕ*. For example, *p <* 1 and *ϕ <* 1 implements a growth-defense trade-off (Fig. 2b).

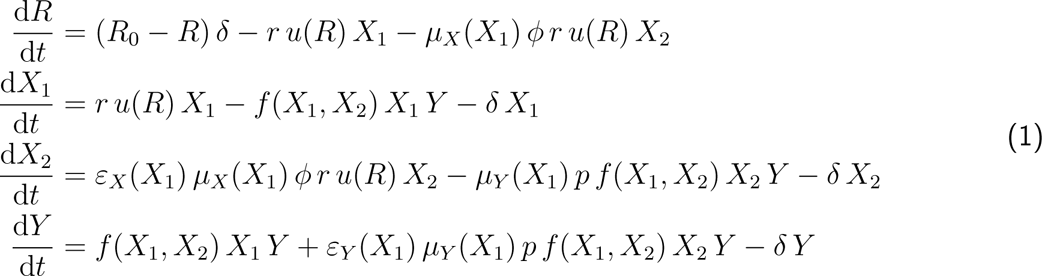

**Figure 1.**
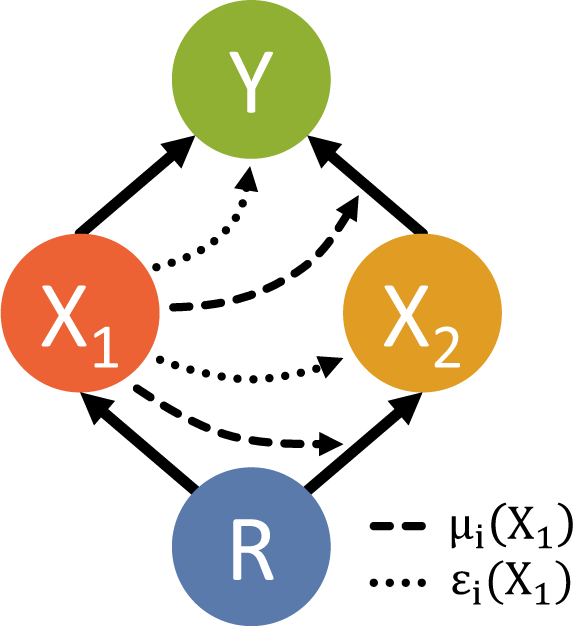
Food web structure. The model equations (Eq. 1) describe a diamond-shaped food web module. Solid arrows depict flows of matter due to resource or prey uptake. Dashed arrows show the interaction modification *µ_i_*(*X*_1_) of the uptake rates that are caused by the provision of essential resources by the focal prey. The other potential higher-order interaction from essential resource provision *ε_i_*(*X*_1_) affects the conversion efficiency of the competitor or the predator and is depicted by dotted arrows. Only one of these higher-order inaction is investigated at a time.

**Figure 2.**
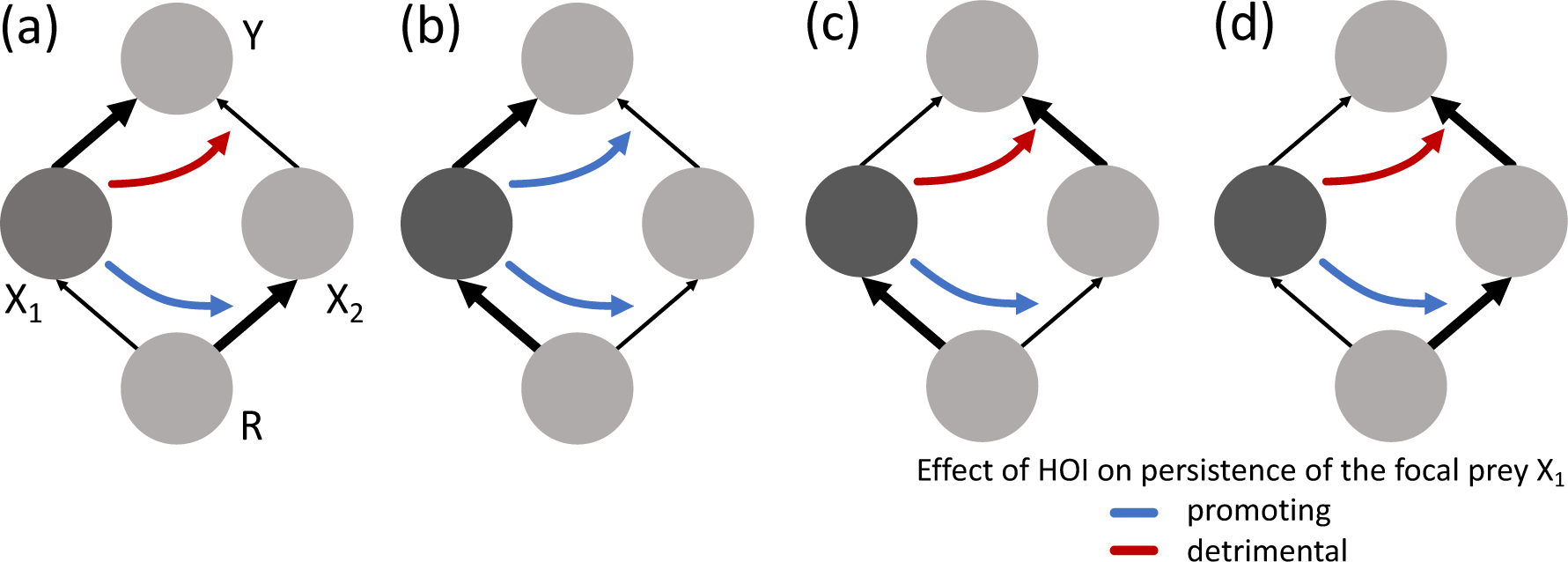
The effect of higher-order interactions depends on the food web scenario. In the first food web scenario (a), the focal prey *X*_1_ is more vulnerable to predation and less competitive than its competitor *X*_2_, whereas it is more vulnerable to predation but also more competitive in the second food web scenario (b). The third (c) and fourth (d) food web scenarios are mirror images of the first and second food web scenario. Essentiality-mediated higher-order interactions that limit the growth of the competitor should favour persistence of the focal prey *X*_1_ (blue curved arrows). Essentiality should promote persistence of the focal prey in food webs that permit predator-mediated coexistence (blue curved arrow in (b)), but likely is detrimental otherwise (red curved arrows) as it can render the competitor effectively less vulnerable to predation than the focal prey.

**Table 1.** Reference parameter set. Resource concentrations and organism abundances or densities are assumed to be normalized appropriately such that their units become unity. Deviations from the reference parameter values are reported where applicable. For an illustration of the different food web scenarios see Fig. 2.

| Parameter | Biological meaning | Value |  |  |  |
| --- | --- | --- | --- | --- | --- |
| $R_0$ | Input concentration of abiotic resource | 1 | | | |
| $\delta$ | Chemostat dilution rate | 1 time unit<sup>-1</sup> | | | |
| $r$ | Prey's uptake rate | 2 time unit<sup>-1</sup> | | | |
| $K$ | Prey's half-saturation constant | 0.1 | | | |
| $g$ | Predator's consumption rate | 1.5 time unit<sup>-1</sup> | | | |
| $H$ | Predator's half-saturation constant | 0.1 | | | |
|  |  | Food web scenarios |  |  |  |
|  |  | I | II | III | IV |
| $\phi$ | Relative competitiveness of the competitor | 1.05 | 0.95 | 0.95 | 1.05 |
| $p$ | Relative vulnerability of the competitor to predation | 0.8 | 0.8 | 1.2 | 1.2 |

Throughout this paper, I use a Monod-type term to indicate resource limitation of the prey

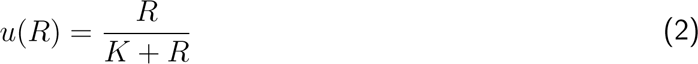

and a Holling Type-2 functional response for multiple prey species to describe the predation rate by an individual predator:

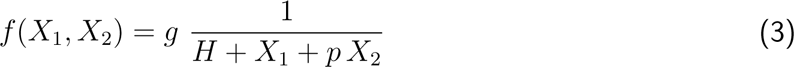

I introduce the higher-order interactions due to essential resource provision as interaction modifications *µ_i_*(*X*_1_) and *ε_i_*(*X*_1_) driven by the density of the focal prey species (Arditi et al., 2005). For generality, I include all possible options where these modifications affect the uptake rates of abiotic resources or prey, or the efficiency at which new biomass is produced, respectively. Accordingly, *µ_X_*(*X*_1_) defines how an increasing density of the focal prey increases the resource uptake rate of the competing prey and *ε_X_*(*X*_1_) gives the conversion efficiency of those resources into new competitor biomass depending on the density of the focal prey. The same logic translates to *µ_Y_* (*X*_1_) and *ε_Y_* (*X*_1_) for the predator, i.e. the provision of essential resources may increase the predation rate on the competitor, for example by alleviating a predator dispreference for the competitor. Similarly, the predator conversion efficiency for competitor biomass may increase due to the provision of essential resources by the focal prey. Note that in this model, I am investigating only the provision of essential resources, thus assuming that the focal prey itself always contains the essential resources. Predator consumption and conversion of focal prey biomass is thus kept constant. I assume that the modification functions *µ_i_*(*X*_1_) and *ε_i_*(*X*_1_) monotonically increase with focal prey density, eventually approaching unity for large focal prey densities, as here their effect should vanish, as the essential resource should be abundantly present and thus non-limiting.

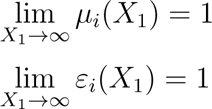

The interaction modifications pose an implicit way of representing the temporal dynamics of production, distribution, stability, uptake and usage of the essential resource molecules and thus avoid the difficulties involved in modelling these processes explicitly, but explicit approaches also exist (Sun et al., 2019; Hammarlund et al., 2019).

I define essentiality *e* as the relative reduction of uptake rates or conversion efficiencies in the absence of the focal prey compared to when it’s abundantly present and neither the uptake rates nor the conversion efficiencies are reduced. For the uptake rate modifications *µ_i_*(*X*_1_) this results in

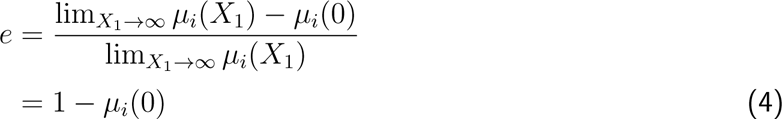

A high essentiality thus implies a strong reduction in the uptake rates when the focal prey is absent.

Similarly, if the higher-order interaction is incorporated into the conversion efficiencies I define

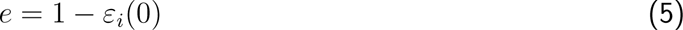

Note that for the sake of simplicity I investigate only the effect of one higher-order interaction at a time, i.e. only one interaction modification will be dependent on the focal prey density, keeping the other three constant at unity.

To determine the effect of essential resource provision on persistence of the focal prey I employ invasion analysis and study whether the focal prey can re-invade the resident community once it has gone extinct. This is ensured by a positive invasion growth rate which is defined as the average per-capita growth rate when rare (Ellner et al., 2019). Specifically, the invasion growth rate of the focal prey in my model is

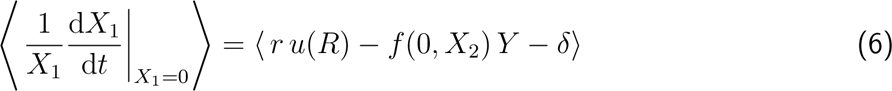

where the angle brackets denote the temporal average. If the resident community’s attractor is a limit cycle, the temporal average can be obtained numerically from one period of such cycles (Ehrlich et al., 2017). As at least some of the parameter combinations investigated in this paper result in limit cycles I used this numerical approach throughout and confirmed the results with the analytically computable solutions for those cases where the resident community was in a steady state. For numerically determining the invasion growth rate of the focal prey, the resident community dynamics were numerically integrated for 2000 time units until they reached their attractor. Convergence was determined visually and verified by ensuring that the absolute values of the slope of the linear regression on the predator abundance, as well as the slope of its moving variance was below 10^−3^ per time unit. The period length was determined as in Raatz et al. (2019) by determining the average time spans between predator maxima during the last 200 time units using the FindMaximum algorithm in Mathematica. The average of the invasion growth rates for each time step during one period was computed. All computations were performed in Mathematica (Wolfram Research, Inc., 2019) and can be re-run using the provided Mathematica notebooks (DOI 10.5281/zenodo.7575588). The analytical solutions are lengthy and can also be found in the notebook and the corresponding pdf exports.

For evaluating the state of the resident community as well as the invasion growth rate of the focal prey, the interaction modifications *µ_i_*(*X*_1_) and *ε_i_*(*X*_1_) reduce to 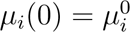 and 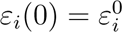, which I can relate to the essentiality *e* via Eqs. 4 and 5. Therefore, I obtain direct relationships between the essentiality of the resource that is provided by the focal prey and its invasion growth rate.

Notably, the invasion analysis does not require a specific choice of the functional form of the interaction modifications. Only those numerical integrations where the focal prey is not set to zero require a particular definition. In those cases I use the following functions that monotonically increase and saturate at unity for large *X*_1_.

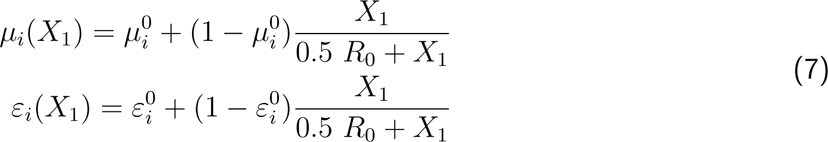

To investigate under which conditions the provision of essential resources can ensure persistence I will focus on four food web scenarios that account for the non-trivial coexistence outcomes in the diamond-shaped food web module. In the first food web scenario the focal prey is the inferior competitor for resource *R* and is more vulnerable to predation than its competitor, which would imply extinction of the focal prey without essentiality (Fig. 2a, see black arrows). In the second food web scenario the focal prey is again more vulnerable to predation but now the superior competitor for resource *R*, which allows for predator-mediated coexistence for a subset of the parameter space, but focal prey extinction otherwise (Abrams, 1999; Jones & Ellner, 2007) (Fig. 2b). The third and fourth food web scenarios are the mirror images of scenarios one and two (Fig. 2c and d). Complementing these scenarios, I will scan the parameter space of vulnerability to predation *p* and resource competitiveness *ϕ* of the competitor relative to the focal prey.

A priori one would expect that essentiality that limits the growth and competitiveness of the competitor should favour the persistence of the focal prey. Further, I hypothesize that within predator-mediated coexistence an increasing essentiality should make the focal prey more indispensable to the community and therefore increase its invasion growth rate, possibly even eventually fulfilling the invasion criterion

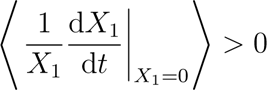

that would prove an ensured persistence of the focal prey.

## Results

Scanning the parameter space of vulnerability to predation *p*and resource competitiveness *ϕ*of the competitor relative to the focal prey provides an overview of the effects of essentiality on the persistence of the focal prey (Fig. 3). Comparing the invasion growth rates at vanishing and complete essentiality, I find that depending on these parameters, and thus the respective food web scenario, essentiality-mediated higher-order interactions can promote but also counter-act the persistence of the focal prey, or have no effect as the focal prey persists or goes extinct irrespective of its essentiality. Analysing the four food web scenarios in more detail provides a detailed understanding of the mechanisms behind these patterns.

**Figure 3.**
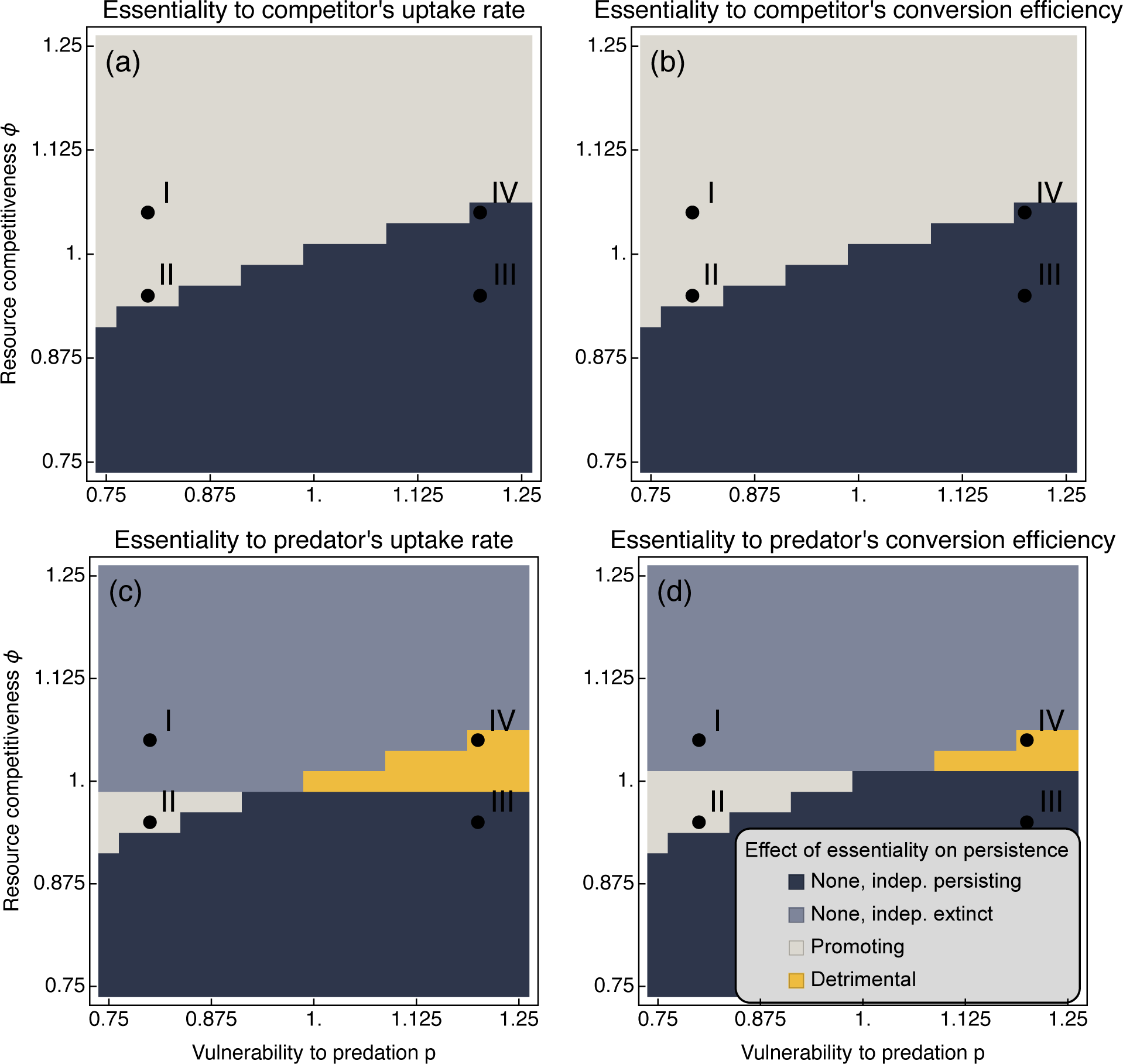
Effect of essentiality on the persistence of the focal prey. Comparing the invasion growth rates of the focal prey for essentialities of *e* = 0 and *e* = 1 allows to classify the effects of essentiality on persistence for the four higher-order interactions indicated in Fig. 1. If the invasion growth rate for vanishing and complete essentiality are both positive then the focal prey persists independent of its essentiality (dark blue region). Vice versa, if both invasion growth rates are negative the focal prey goes extinct independent of its essentiality (light blue region). Sign changes from negative to positive for increasing essentiality indicate a persistence-promoting effect of essentiality (light-grey region), whereas sign changes from positive to negative depict a detrimental effect of essentiality on persistence of the focal prey (yellow region). The parameters of the four food web scenarios of Fig. 2 are indicated by Roman numerals.

In the first food web scenario (Fig. 2a) the focal prey does not persist for vanishing essentiality, as indicated by a negative invasion growth rate. However, increasing essentiality when the higher-order interaction affects the resource uptake rate or conversion efficiency of the competitor turns the invasion growth rate positive (Fig. 4a,b) and thus ensures the persistence of the focal prey (Fig. 5). This includes a drastic shift in the resident community shortly beyond *e* = 0.4 where first the predator and then the prey go extinct (Fig. A1a,b). An essentiality of *e* = 0.4 implies that the resource uptake rate or the conversion efficiency of the competitor are reduced to 60% in the absence of the focal prey. In my model formulation this implies that the competitor cannot sustain the predator further which, in the absence of the focal prey, therefore goes extinct. A slight additional reduction hinders the competitor from outgrowing dilution and thus drives it to extinction as well. In this food web scenario, higher-order interactions that target the uptake rate or the conversion efficiency of the predator do not benefit the persistence of the focal prey (Fig. 4c,d) due to unfavourable trait combinations. As the focal prey is the inferior competitor for the resource *R* and also more vulnerable to predation it can persist neither in the absence nor in the presence of the predator. Supporting the predator by providing essential resources harms the focal prey more than the competitor. For the predator, a larger dependence on the focal prey is also disadvantageous as this decreases its uptake rate and conversion efficiency, and results in extinction at approximately *e* = 0.25 (Fig. A1c,d).

**Figure 4.**
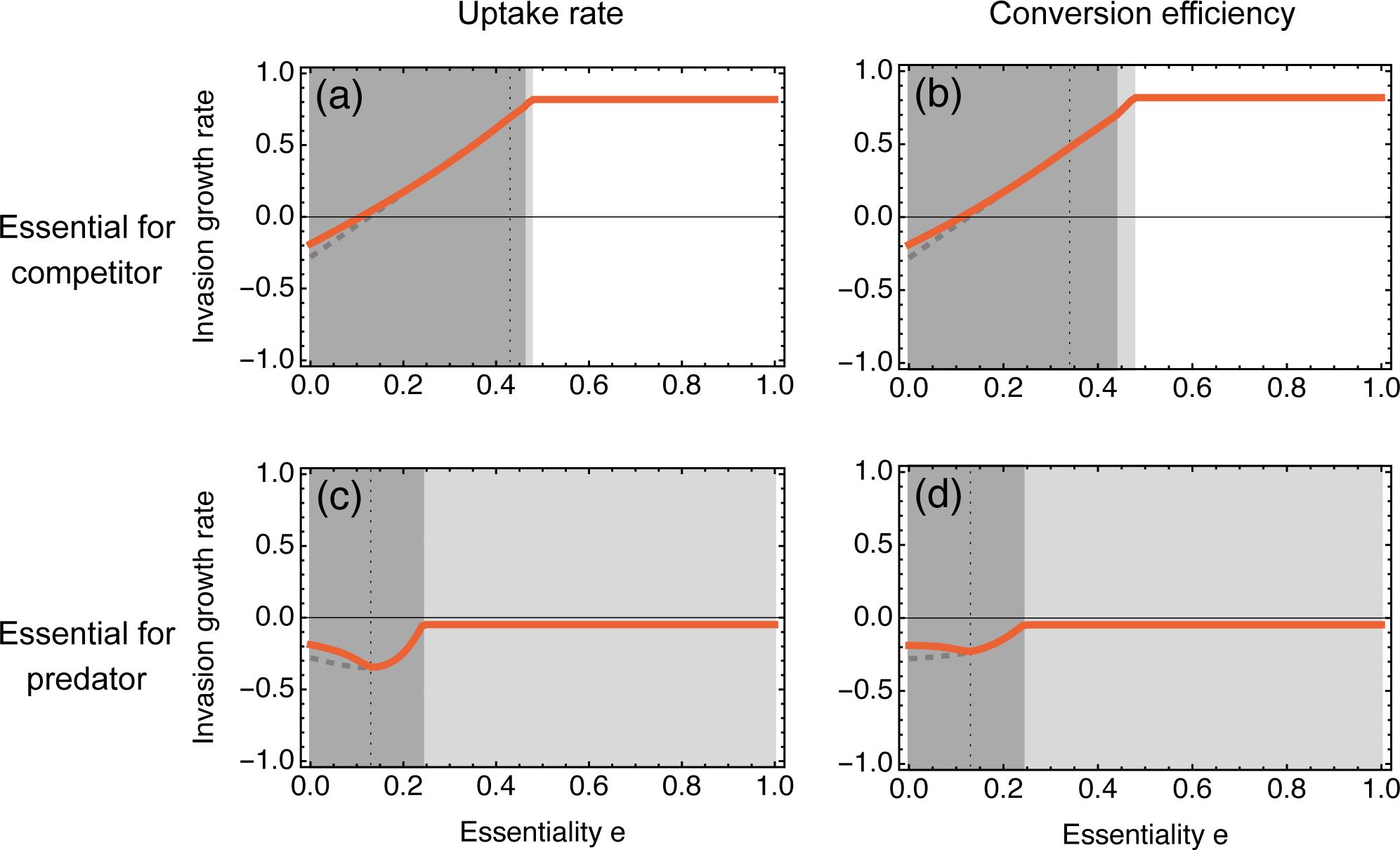
Invasion growth rate of the focal prey for the first food web scenario (Fig. 2a). Here, the focal prey is more vulnerable to predation and competitively inferior to the competitor. Essential resource provisioning affects (a) the uptake rate or (b) the conversion efficiency of the competitor, or (c) the uptake rate or (d) the conversion efficiency of the shared predator. The grey shading indicates the states of the resident community (Fig. A1). For dark shading both predator and competitor coexist, for light-grey shading only the competitor persists and for no shading only the resource remains. The analytically computed invasion growth rate (dashed line) deviates from the numerical observation (full line) once the dynamics become cyclic. The vertical dotted line marks the bifurcation point.

**Figure 5.**
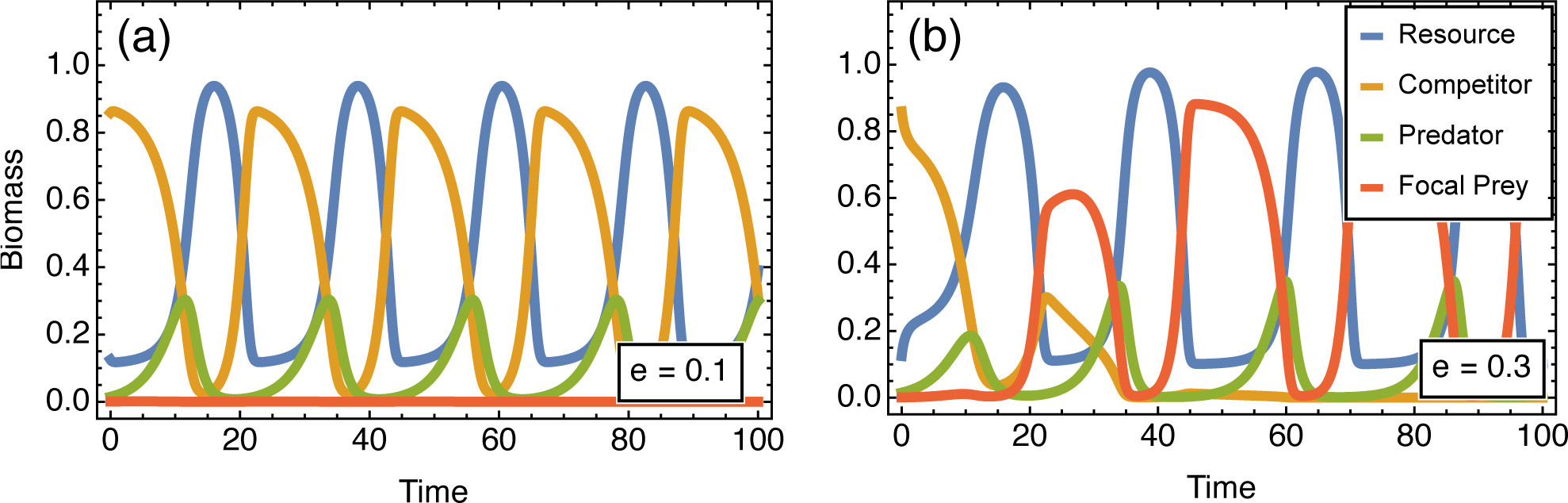
Population dynamics for the first food web scenario when the higher-order interaction targets the resource uptake rate of the competitor (Fig. 4a). (a) For small essentialities the invasion rate of the focal prey is negative and it thus cannot invade. (b) A larger essentiality ensures the persistence of the focal prey. To obtain these dynamics, I chose the interaction modification according to Eq. 7, integrated the resident community to its stable state and then introduced the focal prey at an initial biomass of *X*_1,0_ = 10^−3^.

In the second food web scenario, the focal prey is still more vulnerable to predation than its competitor but now it is also the superior competitor for the resource *R* (Fig. 3). While the invasion analysis outcomes are similar to the first food web scenario for higher-order interactions targeting the competitor’s uptake rate or conversion efficiency (Fig. 6a,b), the trait combinations now allow for positive invasion growth rates also when the higher-order interaction targets the predator’s up-take rate or conversion efficiency (Fig. 6c,d). Therefore, increasing essentiality can promote the persistence of the focal prey for intermediate to large essentiality in this food web scenario. This persistence-promoting effect of essentiality appears in a parameter range of predator-mediated coexistence of prey (Fig. 7). Here, the predator goes extinct in the resident community as the competing prey alone does not sustain the predator given the reduction in uptake rate or conversion efficiency for large essentiality of the focal prey (Fig. A2). In the absence of the predator the focal prey benefits from its higher competitiveness for the resource *R* and thus persists. Once it invades it may additionally sustain the predator (Fig. 7b). Conditional on the presence or absence of the predator when the focal prey invades two community states are therefore possible. Without the predator the focal prey out-competes the competitor which thus goes extinct (Fig. 7b, solid lines). If the predator is present or is re-introduced it however mediates coexistence of the focal prey and the competitor (Fig. 7b, dashed lines). This shows that providing essential resources can affect not only the focal prey itself, but also the whole community structure.

**Figure 6.**
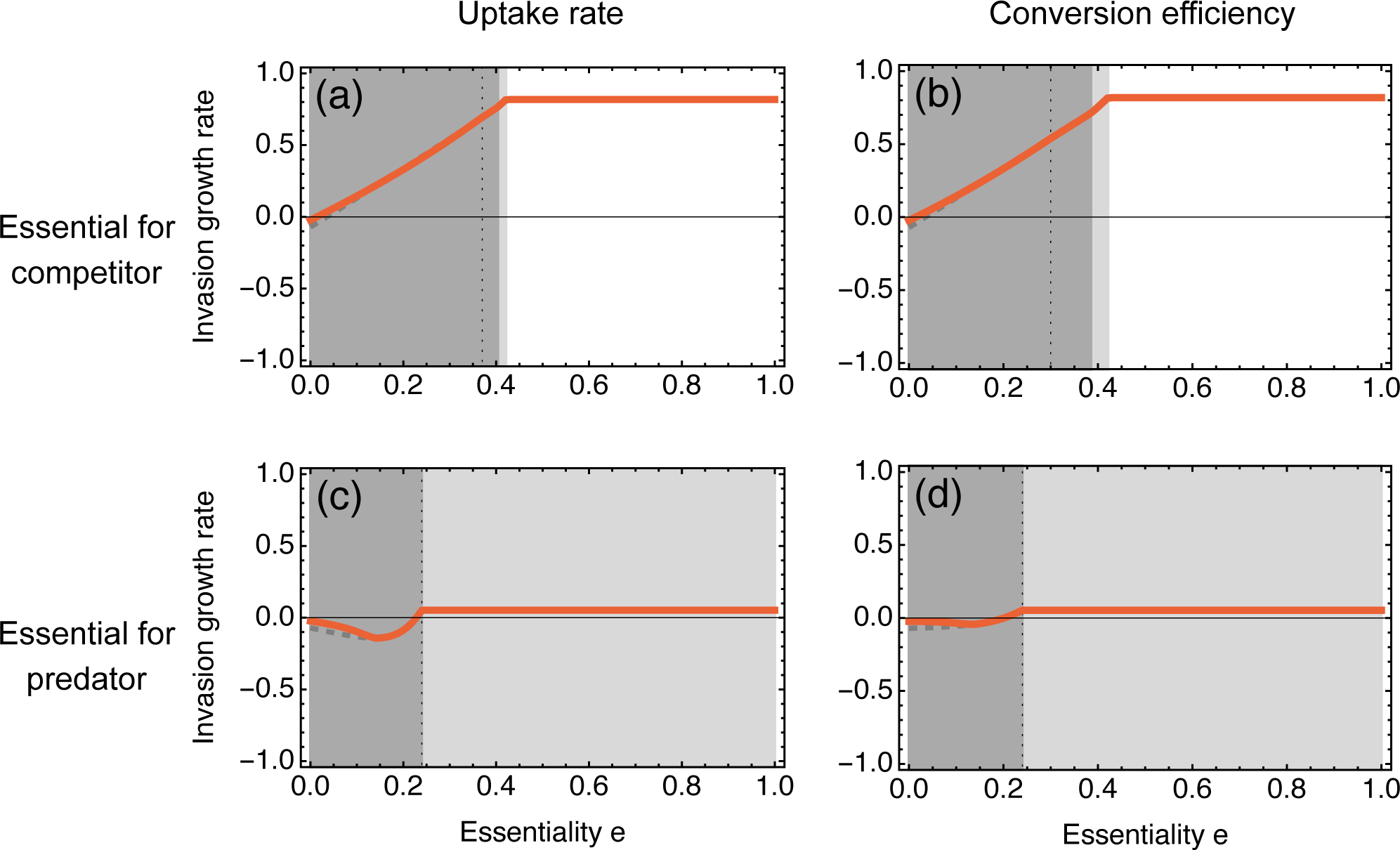
Invasion growth rate for the second food web scenario. Here, the focal prey is more vulnerable to predation but also competitively superior to the competitor. In the absence of the focal prey its essentiality determines the reduction in (a) the uptake rate or (b) the conversion efficiency of the competitor, or (c) the uptake rate or (d) the conversion efficiency of the shared predator. The plot specifics are identical to Fig. 4.

**Figure 7.**
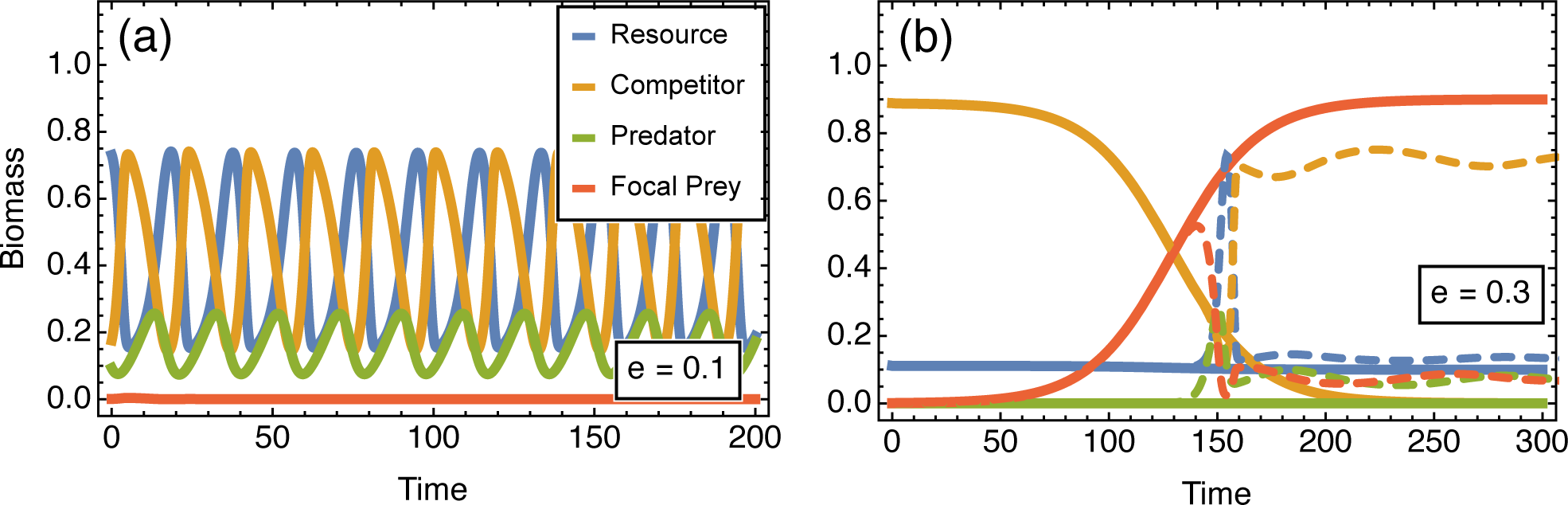
Population dynamics for the second food web scenario when the higher-order interaction targets the resource uptake rate of the shared predator (Fig. 6c). (a) For small essentialities the invasion rate of the focal prey is negative and it thus cannot invade. (b) A larger essentiality ensures the persistence of the focal prey. To obtain these dynamics, I chose the interaction modification according to Eq. 7, integrated the resident community to its stable state and then introduced the focal prey at an initial biomass of *X*_1,0_ = 10^−3^. In panel (b) the predator goes extinct in the residence community, thus I assumed *Y*_0_ = 0 (thick lines). If, however, the predator is reintroduced together with the focal prey (*Y*_0_ = 10^−3^, thin dashed lines), it is supported by the focal prey, re-establishes and mediates the coexistence of both prey types.

In the third food web scenario, the focal prey persists independent of essentiality as indicated by a positive invasion growth rate for all possible types of essentiality-mediated higher order interactions (Fig. 3 and Fig. A3). If essentiality affects the competitor the focal prey’s invasion growth rate increases further. If, however, essentiality causes limitations for the predator the invasion growth rate tends to decrease for larger essentiality (albeit not turning negative) as this effectively reduces the energy flow from the competitor to the predator and thus eventually renders the competitor less vulnerable to predation than the focal prey.

Similarly, higher-order interactions affecting the competitor increase the invasion growth rate of the focal prey with higher essentiality in the fourth food web scenario. If the resource competitiveness of the competitor is only slightly exceeding the resource competitiveness of the focal prey the invasion growth rate of the focal prey is positive even for zero essentiality and only increases further for higher essentiality (Fig. A4). For higher competitiveness of the competitor the invasion growth rate at zero essentiality is negative and turns positive for higher essentiality, again resulting in promoted persistence already (Fig. 3a,b). In this scenario, however, higher-order interactions affecting the predator can result in a negative invasion growth rate, which can become even smaller if the essentiality becomes larger. Here again, an increasing essentiality counteracts the larger vulnerability of the competitor to predation, and allows the competitor to outcompete the focal prey given its higher resource competitiveness.

## Discussion

Higher-order interactions have the potential to shape community structure and dynamics (Grilli et al., 2017; Mayfield & Stouffer, 2017; Terry et al., 2019). In this paper, I showed how the provision of essential resources creates a higher-order interaction that decisively affects the persistence of the focal prey and the resulting community structure. I investigated both the case of essential resource provision to community members from the same trophic level as well as from a higher trophic level. Whether these higher-order interactions in the end ensure persistence depends both on their strength as well as on the food web scenario (see Fig. 2 for a summary of the results).

Confirming the expectations, I find in all food web scenarios that a larger essentiality for the competitor can increase the invasion growth rate of the focal prey. In the first and second food web scenario where the invasion growth rate is negative for zero essentiality this leads to a sign-change in the invasion growth rate and thus a promoting effect of essentiality on persistence. In the third and fourth food web scenarios the invasion growth rate of the focal prey is already positive for zero essentiality and only increases further for larger essentiality. Essentiality for the predator can indeed favour the persistence of the focal prey in food web scenarios that permit predator-mediated coexistence of the prey species (second food web scenario), but can also be detrimental for persistence if it renders the competitor effectively less vulnerable to predation (fourth food web scenario). Further, I find that essentiality determines the resident community structure, with larger essentiality driving extinct first the predator and then, depending on the higher-order interaction, potentially also the competitor. As seen in the second food web scenario this allows for multiple possible community states, depending on whether the coexistence-mediating predator is re-introduced together with the focal prey. Further, no qualitative differences between higher-order interactions affecting the uptake rate or the conversion efficiency were observed.

Experimental support exists for both higher-order interactions that affect the uptake rate or the conversion efficiency. Essential resources affecting the uptake rate could result from adaptive foraging behaviour, as predicted by nutritional geometry (Raubenheimer & Simpson, 1993; Simpson et al., 2004), selective feeding (Buskey, 1997; Elser et al., 2016; Meunier et al., 2016; Eberl et al., 2020), or changed behaviour due to the provision of essential micronutrients, as recently reported for a nematode feeding on larvae of other nematodes (Akduman et al., 2020). Here, the attack rate of the predatory nematode increased when reared on vitamin B_12_ producing bacteria compared to B_12_ deficient controls. However, feeding rate was not increased in this study, so only the prey’s loss term would be affected by this higher-order interaction when transferring these results to my model. Another possibility would be generally better physiological conditions that increase fitness, as reported for *Daphnia magna* and vitamin B_12_ (Kusari et al., 2017), which could also translate to generally increased activity.

The most direct and intuitive mechanism for a higher-order interaction that affects the conversion efficiency of a consumer via essential resource provision is that those lacking essential nutrients that are halting biomass production are directly provided. This is the case in the above example with *Daphnia magna* and vitamin B_12_ (Keating, 1985), other nutrients like phosphorous (Urabe et al., 2018) or biochemicals (Martin-Creuzburg et al., 2009; Raatz et al., 2017). Similarly, supplementing herbivory with fungivory was found to significantly speed up growth in moth larvae (Eberl et al., 2020). Microbial cross-feeding likely represents the case of higher-order interactions affecting the conversion efficiency of organisms on the same trophic level (D’Souza et al., 2018). In the absence of another carbon source bacteria depend on algal carbon fixation and exudation (Bratbak & Thingstad, 1985; Raatz et al., 2018), which was proposed as the mutualistic trade in return for bacterial vitamin B_12_ provision (Croft et al., 2005) during this type of cross-feeding between different kingdoms.

There has been a long history of investigating the effect of higher-order interactions in small ecological interaction networks, such as trait-mediated indirect interactions (Bolker et al., 2003; Werner & Peacor, 2003) or non-trophic interactions (Kéfi et al., 2012), e.g. facilitation (Gross, 2008). The effect of higher-order interactions on community stability is investigated also in larger networks, both empirical (Gonźalez Gonźalez et al., 2021) and theoretical, randomly sampled ones (e.g. Arditi et al., 2005; Grilli et al., 2017; Gibbs et al., 2023), and innovative approaches of analyzing their effects have been proposed (Golubski et al., 2016). The effect of trait-mediated indirect interactions and higher-order interactions in general have been shown to depend on many specifics, such as network structure and interaction strengths. In my model, a higher essentiality corresponds to a higher strength of the higher-order interaction. I found that depending on the food web scenario, food-quality-provision-mediated higher-order interactions can be both promoting but also detrimental to persistence and thus community stability, a finding that resonates with this overall complexity. Exploring the effect of multiple, simultaneously occurring higher-order interactions presents an interesting avenue for future research.

The provision of essential resources changes the abiotic environment of the competitors or predators via changing the pool of available essential resources. It can be seen as a form of niche construction that is implicitly included via an interaction modification between two biotic food web components (similar to Kylafis & Loreau, 2011; Oña et al., 2021). Obviously, the niches of predator and competitor are impacted directly by the presence of the focal prey. Interestingly, however, this niche construction operates also indirectly in the second food web scenario, as the niche of the focal prey is extended through a feedback loop via predator-mediated coexistence of competitor and focal prey. Bridging theory and experiments on higher-order-interactions is challenging (Levine et al., 2017). I worked out that essentiality, defined as the reduction of uptake rates or conversion efficiencies when the focal prey is absent, is an appropriate measure to determine the effect of the higher-order interaction on the persistence of the focal prey, particularly when using invasion analysis. One of the benefits from this definition is that the density-dependent functional form of the higher-order interactions does not need to be specified, which largely facilitates experimental approaches of measuring the presence and effect of the higher-order interactions. In my analysis I focussed on the persistence of the focal prey. It should be noted that determining coexistence of species, and not only persistence of a focal species, can be complicated by the existence of multiple stable states (e.g. Yamamichi et al., 2014) which constrains the interpretation of invasion growth rates (Grainger et al., 2019).

Measuring higher-order interactions experimentally is difficult, however, some advances have been reported that employ different strategies. A first line of research infers the higher-order interactions statistically from community dynamics data (e.g. Kéfi et al., 2015; Mayfield & Stouffer, 2017). A second, more mechanistic approach aims to disable hypothesized higher-order interactions and compare the outcomes with the non-manipulated scenario. One prominent example is the study by Wootton (1993) where the disguising effect of barnacles for limpets was discovered by removing barnacles partially or completely. Removing the species that initiates the higher-order interaction to quantify the effect of the higher-order interaction however is complicated by other direct and indirect effects that are then also removed, which would lead to false evaluations of the effect size of the higher-order interaction. The essential resource context provides a different way of determining the effect size of higher-order interactions. Experimentally providing the essential resource in excess by supplementation removes its potential to cause higher-order interactions and decreases its essentiality.

This approach has been used in investigations of microbial cross-feeding, such as in Kazamia et al. (2012) and Hammarlund et al. (2019) where supplementation with the essential resource alleviated the dependence on the interaction partner, shifting the coexistence pattern towards the beneficiary of the supplementation. In the context of herbivore limitation by biochemicals, supplementation was used to show the mechanistic basis for the higher-order interaction (Wacker & Martin-Creuzburg, 2012). In a predator-prey context Bayesian inference from population size time series can be used to obtain uptake rates and conversion efficiencies independently (Rosenbaum et al., 2019). Applying the inference for different supplementation levels should allow to disentangle whether the essential resource affects the uptake rate or the conversion efficiency of the predator. This may be less feasible for a prey consuming abiotic resources, but even here methods such as isotopic labelling could be used to track uptake and conversion separately. The community-structuring effect of essential resource provision remains to be tested, which requires tracking the population feedback mechanisms over larger time scales of many prey generations, but chemostat or mesocosm experiments will be useful here. The central focus of this article on persistence of the focal prey, however, facilitates experimental validation. As argued before, only the invasion growth rate of the focal prey would have to be obtained for different levels of supplementation with potentially different resident communities. This reduces the time that experimental cultures would have to be operated and avoids experimental difficulties often entailed by long-term observations, ultimately illuminating the potential effect of essential resource provision on prey persistence.

## Acknowledgements

The author would like to thank Elias Ehrlich for helpful comments on an earlier draft. Preprint version 3 of this article has been peer-reviewed and recommended by Peer Community In Ecology. Funding for this research was provided by Max Planck Society.

## Competing Interests

The author states no competing financial interests.

## Data, script and code availability

The Mathematica scripts used for the calculations and for creating the figures have been deposited both as Mathematica notebooks and also as pdf exports in a zenodo repository with DOI 10.5281/zenodo.7575588.

## Appendix

### Supporting figures

**Figure A1.**
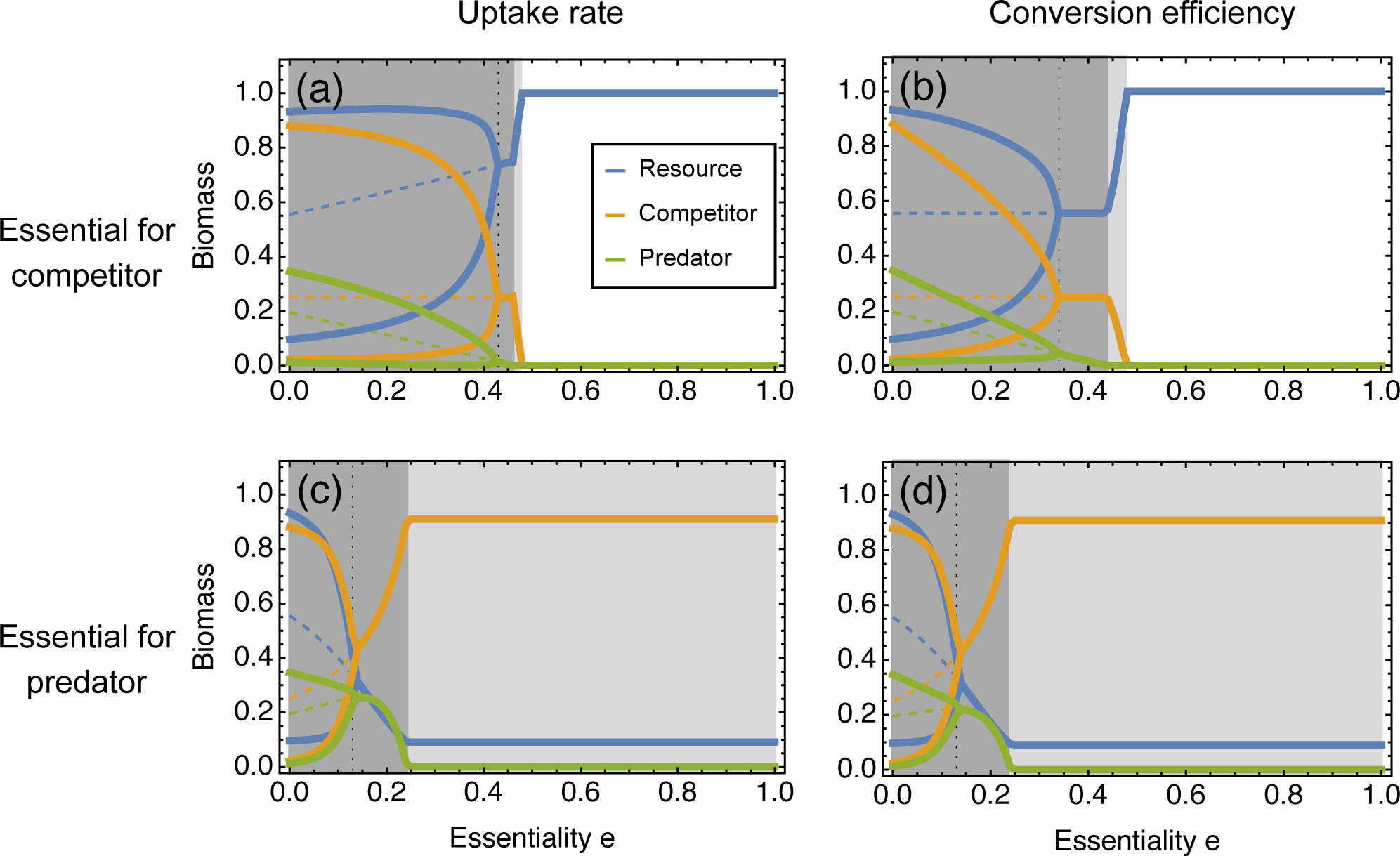
Resident community states for the first food web scenario. In the absence of the focal prey its essentiality determines the reduction in (a) the uptake rate or (b) the conversion efficiency of the competitor, or (c) the uptake rate or (d) the conversion efficiency of the shared predator, which affects their community dynamics. Full lines represent the minima and maxima of one population cycle, if the population is cycling, or otherwise the steady state biomass. The vertical dotted line indicates the bifurcation point. Population dynamics were defined as cyclic if the difference between predator extrema exceeded 10^−5^. During cycles, the unstable fixed point is indicated by the dashed line. As in Fig. 4, the grey shading indicates the states of the resident community. For dark shading both predator and competitor coexist, for light-grey shading only the competitor persists and for no shading only the resource remains.

**Figure A2.**
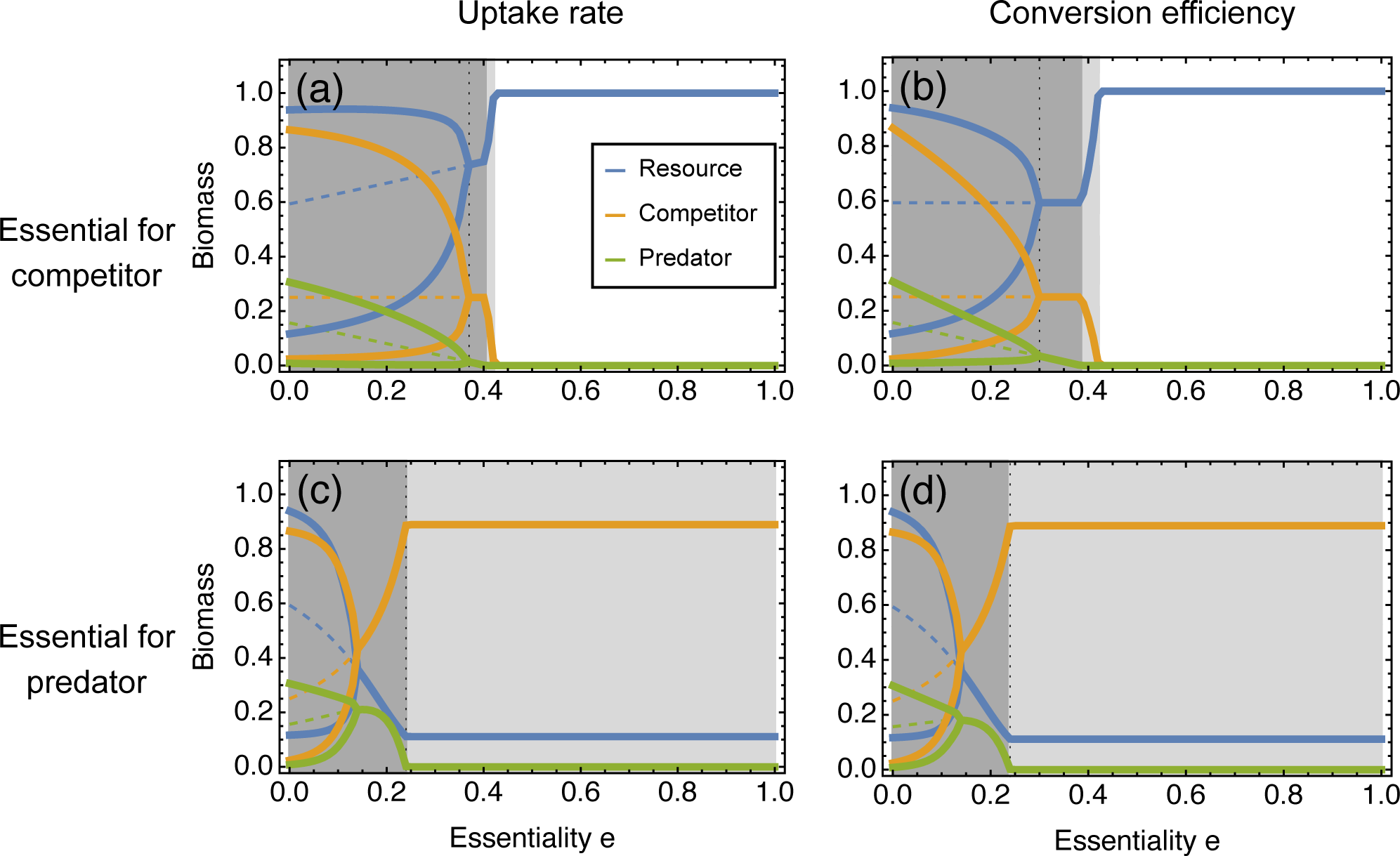
Resident community states for the second food web scenario. Here, the focal prey is more vulnerable to predation but also the superior competitor for the abiotic resource. In the absence of the focal prey its essentiality determines the reduction in (a) the uptake rate or (b) the conversion efficiency of the competitor, or (c) the uptake rate or (d) the conversion efficiency of the shared predator. The plot specifics are identical to Fig. A1.

**Figure A3.**
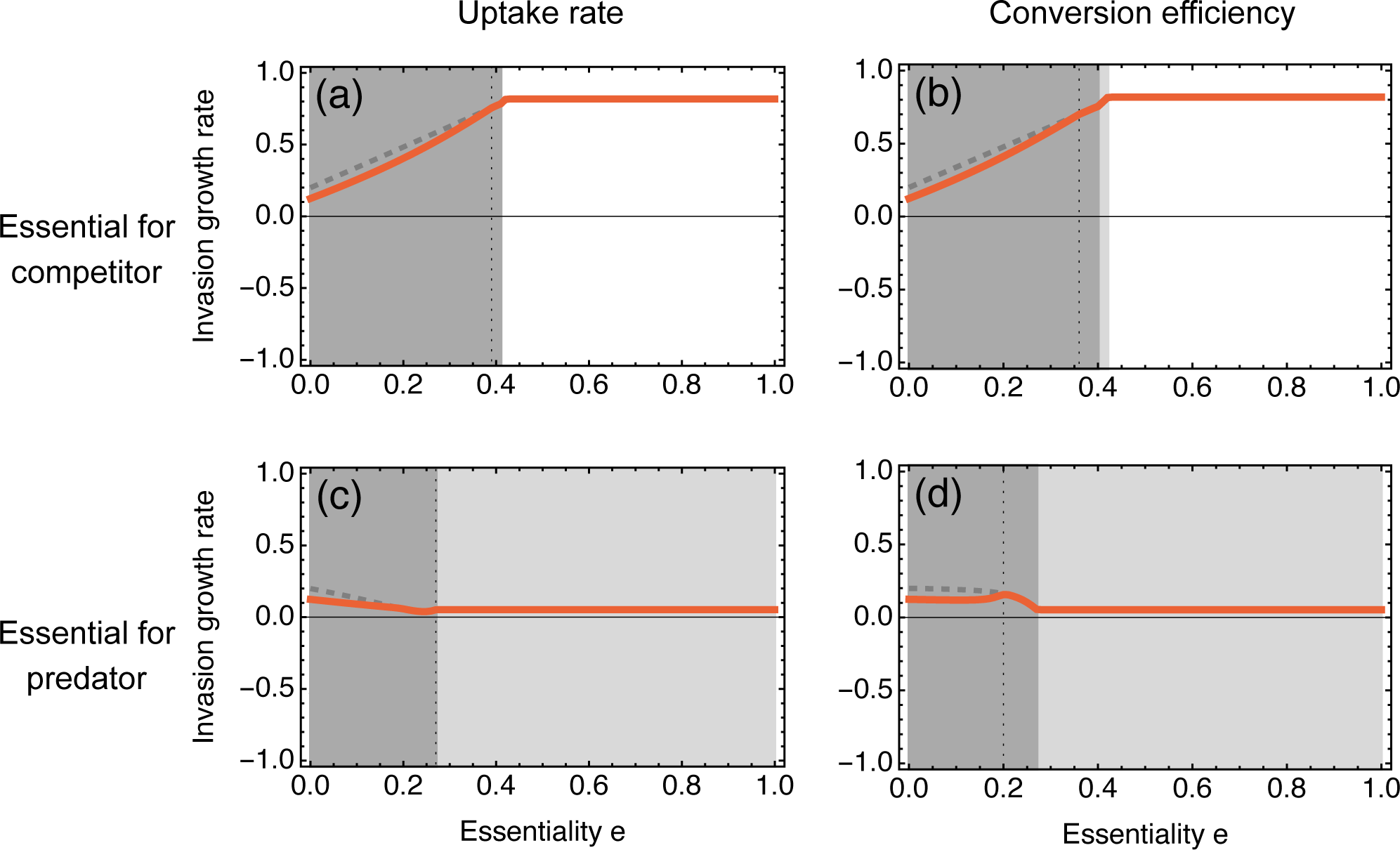
Invasion growth rate of the focal prey for the third food web scenario (Fig. 2c). Here, the focal prey is less vulnerable to predation and more competitive for the resource than the competitor. Essential resource provisioning affects (a) the uptake rate or (b) the conversion efficiency of the competitor, or (c) the uptake rate or (d) the conversion efficiency of the shared predator. Further plot specifics are identical to Figs. 4 and 6.

**Figure A4.**
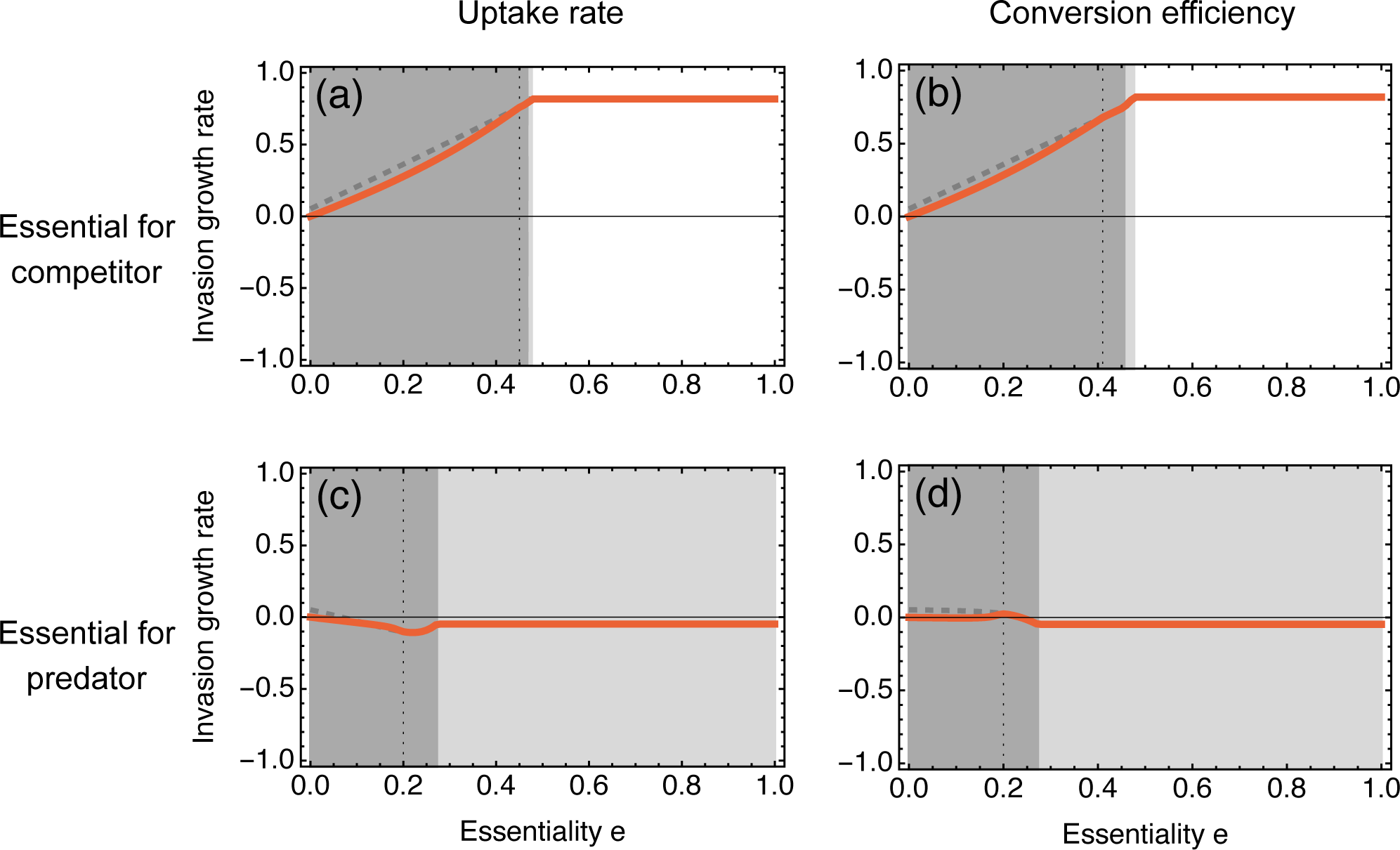
Invasion growth rate of the focal prey for the fourth food web scenario (Fig. 2d). Here, the focal prey is less vulnerable to predation and less competitive for the resource than the competitor. Essential resource provisioning affects (a) the uptake rate or (b) the conversion efficiency of the competitor, or (c) the uptake rate or (d) the conversion efficiency of the shared predator. Further plot specifics are identical to Figs. 4 and 6.

## Notes

https://doi.org/10.5281/zenodo.7575588

### Competing Interest Statement

The authors have declared no competing interest.

### Summary of Updates

PCI Ecology badge added and line numbers removed.

## References

Abrams, P. A. (1999). Is Predator-Mediated Coexistence Possible in Unstable Systems? Ecology, 80(2), 608–621. 10.2307/176639

Adamowicz, E. M., Flynn, J., Hunter, R. C., & Harcombe, W. R. (2018). Cross-feeding modulates antibiotic tolerance in bacterial communities. The ISME Journal, 12(11), 2723–2735. 10.1038/s41396-018-0212-z

Akduman, N., Lightfoot, J. W., Röseler, W., Witte, H., Lo, W.-S., Rödelsperger, C., & Sommer, R. J. (2020). Bacterial vitamin B12 production enhances nematode predatory behavior. The ISME Journal, 14(6), 1494–1507. 10.1038/s41396-020-0626-2

Andersen, T., Elser, J. J., & Hessen, D. O. (2004). Stoichiometry and population dynamics: Stoichiometry and population dynamics. Ecology Letters, 7(9), 884–900. 10.1111/j.1461-0248.2004.00646.x

Anderson, T. R. & Hessen, D. O. (2005). Threshold elemental ratios for carbon versus phosphorus limitation in Daphnia. Freshwater Biology, 50(12), 2063–2075. 10.1111/j.1365-2427.2005.01450.x

Arditi, R., Michalski, J., & Hirzel, A. H. (2005). Rheagogies: Modelling non-trophic effects in food webs. Ecological Complexity, 2(3), 249–258. 10.1016/j.ecocom.2005.04.003

Azam, F., Fenchel, T., Field, JG., Gray, JS., Meyer-Reil, LA., & Thingstad, F. (1983). The Ecological Role of Water-Column Microbes in the Sea. Marine Ecology Progress Series, 10(November 2015), 257–263. 10.3354/meps010257

Billick, I. & Case, T. J. (1994). Higher Order Interactions in Ecological Communities: What Are They and How Can They be Detected? Ecology, 75(6), 1529–1543. 10.2307/1939614

Bolker, B., Holyoak, M., Ǩrivan, V., Rowe, L., & Schmitz, O. (2003). Connecting Theoretical and Empirical Studies of Trait-Mediated Interactions. Ecology, 84(5), 1101–1114. 10.1890/0012-9658(2003)084[1101:CTAESO]2.0.CO;2

Bratbak, G. & Thingstad, T. F. (1985). Phytoplankton-bacteria interactions: An apparent paradox? Analysis of a model system with both competition and commensalism. Marine Ecology Progress Series, 25, 23–30. 10.1016/0198-0254(86)91170-2

Burian, A., Nielsen, J. M., & Winder, M. (2020). Food quantity–quality interactions and their impact on consumer behavior and trophic transfer. Ecological Monographs, 90(1). 10.1002/ecm.1395

Buskey, E. J. (1997). Behavioral components of feeding selectivity of the heterotrophic dineflagellate Protoperidinium pellucidum. Marine Ecology Progress Series, 153(1-3), 77–89. 10.3354/meps153077

Chesson, P. (1994). Multispecies Competition in Variable Environments. Theoretical Population Biology, 45(3), 227–276. 10.1006/tpbi.1994.1013

Croft, M. T., Lawrence, A. D., Raux-Deery, E., Warren, M. J., & Smith, A. G. (2005). Algae acquire vitamin B12 through a symbiotic relationship with bacteria. Nature, 438(7064), 90–93. 10.1038/nature04056

Douglas, A. E. (2015). Multiorganismal Insects: Diversity and Function of Resident Microorganisms. Annual Review of Entomology, 60(1), 17–34. 10.1146/annurev-ento-010814-020822

D’Souza, G., Shitut, S., Preussger, D., Yousif, G., Waschina, S., & Kost, C. (2018). Ecology and evolution of metabolic cross-feeding interactions in bacteria. Natural Product Reports, 35(5), 455–488. 10.1039/c8np00009c

Eberl, F., Fernandez de Bobadilla, M., Reichelt, M., Hammerbacher, A., Gershenzon, J., & Unsicker, S. B. (2020). Herbivory meets fungivory: Insect herbivores feed on plant pathogenic fungi for their own benefit. Ecology Letters, 23(7), 1073–1084. 10.1111/ele.13506

Ehrlich, E., Becks, L., & Gaedke, U. (2017). Trait-fitness relationships determine how trade-off shapes affect species coexistence. Ecology, 98(12), 3188–3198. 10.1002/ecy.2047

Ehrlich, E. & Gaedke, U. (2018). Not attackable or not crackable – How pre- and post-attack defenses with different competition costs affect prey coexistence and population dynamics. Ecology and Evolution, 8(13), 6625–6637. 10.1002/ece3.4145

Ellner, S. P., Snyder, R. E., Adler, P. B., & Hooker, G. (2019). An expanded modern coexistence theory for empirical applications. Ecology Letters, 22(1), 3–18. 10.1111/ele.13159

Elser, J., Kyle, M., Learned, J., McCrackin, M., Peace, A., & Steger, L. (2016). Life on the stoichiometric knife-edge: Effects of high and low food C:P ratio on growth, feeding, and respiration in three Daphnia species. Inland Waters, 6(2), 136–146. 10.5268/IW-6.2.908

Elser, J. J., Dobberfuhl, D. R., MacKay, N. A., & Schampel, J. H. (1996). Organism Size, Life History, and N:P Stoichiometry. BioScience, 46(9), 674–684. 10.2307/1312897

Filipiak, M., Kuszewska, K., Asselman, M., Denisow, B., Stawiarz, E., Woyciechowski, M., & Weiner, J. (2017). Ecological stoichiometry of the honeybee: Pollen diversity and adequate species composition are needed to mitigate limitations imposed on the growth and development of bees by pollen quality. PLOS ONE, 12(8), e0183236. 10.1371/journal.pone.0183236

Gaedke, U., Hochstädter, S., & Straile, D. (2002). Interplay between energy limitation and nutritional deficiency: Empirical data and food web models. Ecological Monographs, 72(2), 251–270. 10.1890/0012-9615(2002)072[0251:IBELAN]2.0.CO;2

Gibbs, T., Gellner, G., Levin, S. A., McCann, K. S., Hastings, A., & Levine, J. M. (2023). Can higher-order interactions resolve the species coexistence paradox? bioRxiv. 10.1101/2023.06.19.545649

Golubski, A. J., Westlund, E. E., Vandermeer, J., & Pascual, M. (2016). Ecological Networks over the Edge: Hypergraph Trait-Mediated Indirect Interaction (TMII) Structure. Trends in Ecology & Evolution, 31(5), 344–354. 10.1016/j.tree.2016.02.006

Gonźalez Gonźalez, C., Mora Van Cauwelaert, E., Boyer, D., Perfecto, I., Vandermeer, J., & Beńıtez, M. (2021). High-order interactions maintain or enhance structural robustness of a coffee agroecosystem network. Ecological Complexity, 47, 100951. 10.1016/j.ecocom.2021.100951

Gore, J., Youk, H., & Van Oudenaarden, A. (2009). Snowdrift game dynamics and facultative cheating in yeast. Nature, 459(7244), 253–256. 10.1038/nature07921

Grainger, T. N., Levine, J. M., & Gilbert, B. (2019). The Invasion Criterion: A Common Currency for Ecological Research. Trends in Ecology & Evolution, 34(10), 925–935. 10.1016/j.tree.2019.05.007

Grilli, J., Barabás, G., Michalska-Smith, M. J., & Allesina, S. (2017). Higher-order interactions stabilize dynamics in competitive network models. Nature, 548(7666), 210–213. 10.1038/nature23273

Gross, K. (2008). Positive interactions among competitors can produce species-rich communities. Ecology Letters, 11(9), 929–936. 10.1111/j.1461-0248.2008.01204.x

Guo, F., Kainz, M. J., Sheldon, F., & Bunn, S. E. (2016). The importance of high-quality algal food sources in stream food webs - current status and future perspectives. Freshwater Biology, 61(6), 815–831. 10.1111/fwb.12755

Hammarlund, S. P., Chaćon, J. M., & Harcombe, W. R. (2019). A shared limiting resource leads to competitive exclusion in a cross-feeding system. Environmental Microbiology, 21(2), 759–771. 10.1111/1462-2920.14493

Herren, C. M. (2020). Disruption of cross-feeding interactions by invading taxa can cause invasional meltdown in microbial communities. Proceedings of the Royal Society B: Biological Sciences, 287(1927), 20192945. 10.1098/rspb.2019.2945

Iwabuchi, T. & Urabe, J. (2012). Food quality and food threshold: Implications of food stoichiometry to competitive ability of herbivore plankton. Ecosphere, 3(6), art51–art51. 10.1890/ES12-00098.1

Jones, L. E. & Ellner, S. P. (2007). Effects of rapid prey evolution on predator–prey cycles. Journal of Mathematical Biology, 55(4), 541–573. 10.1007/s00285-007-0094-6

Kazamia, E., Czesnick, H., Nguyen, T. T. V., Croft, M. T., Sherwood, E., Sasso, S., Hodson, S. J., Warren, M. J., & Smith, A. G. (2012). Mutualistic interactions between vitamin B12-dependent algae and heterotrophic bacteria exhibit regulation. Environmental Microbiology, 14(6), 1466–1476. 10.1111/j.1462-2920.2012.02733.x

Keating, K. I. (1985). The Influence of Vitamin B12 Deficiency on the Reproduction of Daphnia Pulex Leydig (Cladocera). Journal of Crustacean Biology, 5(1), 130–136. 10.2307/1548225

Kéfi, S., Berlow, E. L., Wieters, E. A., Joppa, L. N., Wood, S. A., Brose, U., & Navarrete, S. A. (2015). Network structure beyond food webs: Mapping non-trophic and trophic interactions on Chilean rocky shores. Ecology, 96(1), 291–303. 10.1890/13-1424.1

Kéfi, S., Berlow, E. L., Wieters, E. A., Navarrete, S. A., Petchey, O. L., Wood, S. A., Boit, A., Joppa, L. N., Lafferty, K. D., Williams, R. J., Martinez, N. D., Menge, B. A., Blanchette, C. A., Iles, A. C., & Brose, U. (2012). More than a meal… integrating non-feeding interactions into food webs. Ecology Letters, 15(4), 291–300. 10.1111/j.1461-0248.2011.01732.x

Koedooder, C., Stock, W., Willems, A., Mangelinckx, S., De Troch, M., Vyverman, W., & Sabbe, K. (2019). Diatom-Bacteria Interactions Modulate the Composition and Productivity of Benthic Diatom Biofilms. Frontiers in Microbiology, 10. 10.3389/fmicb.2019.01255

Koussoroplis, A. M., Schälicke, S., Raatz, M., Bach, M., & Wacker, A. (2019). Feeding in the frequency domain: Coarser-grained environments increase consumer sensitivity to resource variability, covariance and phase. Ecology Letters, 22(7), 1104–1114. 10.1111/ele.13267

Kusari, F., O’Doherty, A. M., Hodges, N. J., & Wojewodzic, M. W. (2017). Bi-directional effects of vitamin B 12 and methotrexate on Daphnia magna fitness and genomic methylation. Scientific Reports, 7(1), 11872. 10.1038/s41598-017-12148-2

Kylafis, G. & Loreau, M. (2008). Ecological and evolutionary consequences of niche construction for its agent. Ecology Letters, 11(10), 1072–1081. 10.1111/j.1461-0248.2008.01220.x

Kylafis, G. & Loreau, M. (2011). Niche construction in the light of niche theory. Ecology Letters, 14(2), 82–90. 10.1111/j.1461-0248.2010.01551.x

Laland, K., Matthews, B., & Feldman, M. W. (2016). An introduction to niche construction theory. Evolutionary Ecology, 30(2), 191–202. 10.1007/s10682-016-9821-z

Levine, J. M., Bascompte, J., Adler, P. B., & Allesina, S. (2017). Beyond pairwise mechanisms of species coexistence in complex communities. Nature, 546(7656), 56–64. 10.1038/nature22898

MacArthur, R. & Levins, R. (1967). The Limiting Similarity, Convergence, and Divergence of Coexisting Species. The American Naturalist, 101(921), 377–385. 10.1086/282505

Martin-Creuzburg, D., Sperfeld, E., & Wacker, A. (2009). Colimitation of a freshwater herbivore by sterols and polyunsaturated fatty acids. Proceedings of the Royal Society B: Biological Sciences, 276(1663), 1805–1814. 10.1098/rspb.2008.1540

Mayfield, M. M. & Stouffer, D. B. (2017). Higher-order interactions capture unexplained complexity in diverse communities. Nature Ecology and Evolution, 1(3), 1–7. 10.1038/s41559-016-0062

Meunier, C. L., Boersma, M., Wiltshire, K. H., & Malzahn, A. M. (2016). Zooplankton eat what they need: Copepod selective feeding and potential consequences for marine systems. Oikos, 125(1), 50–58. 10.1111/oik.02072

Miele, V., Guill, C., Ramos-Jiliberto, R., & Kéfi, S. (2019). Non-trophic interactions strengthen the diversity—functioning relationship in an ecological bioenergetic network model. PLoS Computational Biology, 15(8). 10.1371/journal.pcbi.1007269

Muller, E. B., Nisbet, R. M., Kooijman, S. A. L. M., Elser, J. J., & McCauley, E. (2001). Stoichiometric food quality and herbivore dynamics. Ecology Letters, 4(6), 519–529. 10.1046/j.1461-0248.2001.00240.x

Oña, L., Giri, S., Avermann, N., Kreienbaum, M., Thormann, K. M., & Kost, C. (2021). Obligate cross-feeding expands the metabolic niche of bacteria. Nature Ecology & Evolution, 5(9), 1224–1232. 10.1038/s41559-021-01505-0

Oña, L. & Kost, C. (2022). Cooperation increases robustness to ecological disturbance in microbial cross-feeding networks. Ecology Letters, 25(6), 1410–1420. 10.1111/ele.14006

Pomeroy, L., leB. Williams, P. J., Azam, F., & Hobbie, J. (2007). The Microbial Loop. Oceanography, 20(2), 28–33. 10.5670/oceanog.2007.45

Raatz, M., Gaedke, U., & Wacker, A. (2017). High food quality of prey lowers its risk of extinction. Oikos, 126(10), 1501–1510. 10.1111/oik.03863

Raatz, M., Schälicke, S., Sieber, M., Wacker, A., & Gaedke, U. (2018). One man’s trash is another man’s treasure-the effect of bacteria on phytoplankton-zooplankton interactions in chemostat systems: Bacteria in chemostat experiments. Limnology and Oceanography: Methods, 16(10), 629–639. 10.1002/lom3.10269

Raatz, M., van Velzen, E., & Gaedke, U. (2019). Co-adaptation impacts the robustness of predator–prey dynamics against perturbations. Ecology and Evolution, 9(7), 3823–3836. 10.1002/ece3.5006

Raubenheimer, D. & Simpson, S. J. (1993). The geometry of compensatory feeding in the locust. Animal Behaviour, 45(5), 953–964. 10.1006/anbe.1993.1114

Rosenbaum, B., Raatz, M., Weithoff, G., Fussmann, G. F., & Gaedke, U. (2019). Estimating Parameters From Multiple Time Series of Population Dynamics Using Bayesian Inference. Frontiers in Ecology and Evolution, 6, 234–234. 10.3389/fevo.2018.00234

Schade, J. D., Kyle, M., Hobbie, S. E., Fagan, W. F., & Elser, J. J. (2003). Stoichiometric tracking of soil nutrients by a desert insect herbivore. Ecology Letters, 6(2), 96–101. 10.1046/j.1461-0248.2003.00409.x

Schälicke, S., Sobisch, L.-Y., Martin-Creuzburg, D., & Wacker, A. (2019). Food quantity–quality colimitation: Interactive effects of dietary carbon and essential lipid supply on population growth of a freshwater rotifer. Freshwater Biology, 64(5), 903–912. 10.1111/fwb.13272

Simpson, S. J., Sibly, R. M., Lee, K. P., Behmer, S. T., & Raubenheimer, D. (2004). Optimal foraging when regulating intake of multiple nutrients. Animal Behaviour, 68(6), 1299–1311. 10.1016/j.anbehav.2004.03.003

Singer, M. S., Farkas, T. E., Skorik, C. M., & a Mooney, K. (2012). Tritrophic interactions at a community level: Effects of host plant species quality on bird predation of caterpillars. The American naturalist, 179(3), 363–374. 10.1086/664080

Soria-Dengg, S., Reissbrodt, R., & Horstmann, U. (2001). Siderophores in marine coastal waters and their relevance for iron uptake by phytoplankton: Experiments with the diatom Phaeodactylum tricornutum. Marine Ecology Progress Series, 220, 73–82. 10.3354/meps220073

Stiefs, D., van Voorn, G. A. K., Kooi, B. W., Feudel, U., & Gross, T. (2010). Food Quality in Producer-Grazer Models: A Generalized Analysis. The American Naturalist, 176(3), 367–380. 10.1086/655429

Suleiman, M., Zecher, K., Yücel, O., Jagmann, N., & Philipp, B. (2016). Interkingdom Cross-Feeding of Ammonium from Marine Methylamine-Degrading Bacteria to the Diatom Phaeodactylum tricornutum. Applied and Environmental Microbiology, 82(24), 7113–7122. 10.1128/AEM.01642-16

Sun, Z., Koffel, T., Stump, S. M., Grimaud, G. M., & Klausmeier, C. A. (2019). Microbial cross-feeding promotes multiple stable states and species coexistence, but also susceptibility to cheaters. Journal of Theoretical Biology, 465, 63–77. 10.1016/j.jtbi.2019.01.009

Terry, J. C. D., Morris, R. J., & Bonsall, M. B. (2019). Interaction modifications lead to greater robustness than pairwise non-trophic effects in food webs. Journal of Animal Ecology, 88(11), 1732–1742. 10.1111/1365-2656.13057

Urabe, J., Shimizu, Y., & Yamaguchi, T. (2018). Understanding the stoichiometric limitation of herbivore growth: The importance of feeding and assimilation flexibilities. Ecology Letters, 21(2), 197–206. 10.1111/ele.12882

Wacker, A. & Martin-Creuzburg, D. (2012). Biochemical nutrient requirements of the rotifer Brachionus calyciflorus: Co-limitation by sterols and amino acids. Functional Ecology, 26(5), 1135–1143. 10.1111/j.1365-2435.2012.02047.x

Werner, E. E. & Peacor, S. D. (2003). A review of trait-mediated indirect interactions in ecological communities. Ecology, 84(5), 1083–1100.

Wolfram Research, Inc. (2019). Mathematica, Version 12.0. Wolfram Research, Inc., Champaign, IL.

Wootton, J. T. (1993). Indirect Effects and Habitat Use in an Intertidal Community: Interaction Chains and Interaction Modifications. The American Naturalist, 141(1), 71–89. 10.1086/285461

Wootton, J. T. (2002). Indirect effects in complex ecosystems: Recent progress and future challenges. Journal of Sea Research, 48(2), 157–172. 10.1016/S1385-1101(02)00149-1

Yamamichi, M., Yoshida, T., & Sasaki, A. (2014). Timing and propagule size of invasion determine its success by a time-varying threshold of demographic regime shift. Ecology, 95(8), 2303–2315. 10.1890/13-1527.1

